# A data-driven approach to selecting invited speakers at conferences: a step toward gender parity

**DOI:** 10.1101/426320

**Authors:** Ann-Maree Vallence, Mark R Hinder, Hakuei Fujiyama

## Abstract

We present a data-driven approach that uses established metrics of scientific quality to select invited speakers; this approach enables gender parity in conference programs while ensuring high scientific standards.

---

Gender disparity in academia has been acknowledged for some time. In science, technology, engineering, and mathematics (STEM), women represent approximately half of PhD graduates but only approximately a quarter of professors (1). Although calls for approaches to help achieve gender parity in STEM have been numerous, progress is slow.

Recently, gender disparity in invited speaker presentations at scientific conferences has attracted much attention, with evidence of such disparity in STEM conference programs including (but not limited to) fields such as sport and exercise medicine (2), evolutionary biology (3), mathematics (4), ecology (5), and microbiology (6). In the field of neuroscience, BiasWatchNeuro has published gender data of speakers to increase accountability for gender disparity in conference programs; data extracted from BiasWatchNeuro (on 12/20/18) indicated that only 27% of invited speakers across ∼400 neuroscience conferences (between 2014-2018) were women (7). Although some neuroscience conferences are attaining, or exceeding, parity in invited speakers, more than 80% of conferences have fewer than 50% women in their invited speaker programs (7). Given that such opportunities are critical for career development, approaches to achieve gender parity in conference programs while maintaining the high scientific standards expected in conferences programs are needed.

In recent years, some neuroscience societies have developed and implemented equity and diversity policies that can guide the composition of conference programs (8) (e.g. Australasian Cognitive Neuroscience Society https://www.acns.org.au/wp-content/uploads/2018/08/ACNS_Equity_Diversity_Policy_Final_Nov2016.pdf). In addition, online calculators can provide estimates of equitable gender representation, and explicit guidelines for conference committees can prescribe equitable gender representation (9). The development and uptake of these approaches provide evidence of a willingness to compose more equitable conferences and improve the conventional subjective speaker selection process. However, none of the aforementioned approaches provide information regarding *who* to invite as conference speakers, and equity and diversity policies are often subject to the criticism that the selection process is not based on scientific quality. The subjective nature of speaker selection decisions, as well as the criticism that policies and quotas are not scientific quality-based, might (partly) underlie the persistent gender disparity in neuroscience conference programs. The current project was conceived to directly address this issue.

We developed a two-step approach to invited speaker selection, aimed at achieving conference programs with high scientific quality, individually identified speakers, and gender parity. First, we audited the top ten neuroscience journals (indexed by SCImago Journal and Country Rank; SJR), identifying (i) highly cited papers, as well as (ii) gender, (iii) field-weighted citation impact (FWCI), and (iv) total publication count of the first and last authors. Second, we used these data to establish a database of high quality scientists from which speakers could be selected for conferences. Establishing such a database of high quality researchers based on these metrics can provide convincing evidence of parity in scientific quality between men and women at the highest level: if the *quality* of men and women on this database is indistinguishable, this approach permits gender balanced invitations that are based on established metrics of quality frequently used by researchers, hiring committees, and funding bodies. That is to say, the approach presented here extends beyond the currently available tools by (i) identifying particular individuals as potential speakers and (ii) overcoming the criticism that selection based on policies and quotas alone is not based on scientific quality. Notably, this approach can have an immediate effect to improve the representation of women invited speakers at neuroscience conferences, and will likely have a medium-to-long-term effect to improve the progression of women scientists to senior levels within STEM.

## Method

The study was approved by the Murdoch University Human Research Ethics Committee (2017/206). Figure 1 shows the study procedure (journal ranking data and citation reports extracted 11/26/2017).

**Figure 1.**
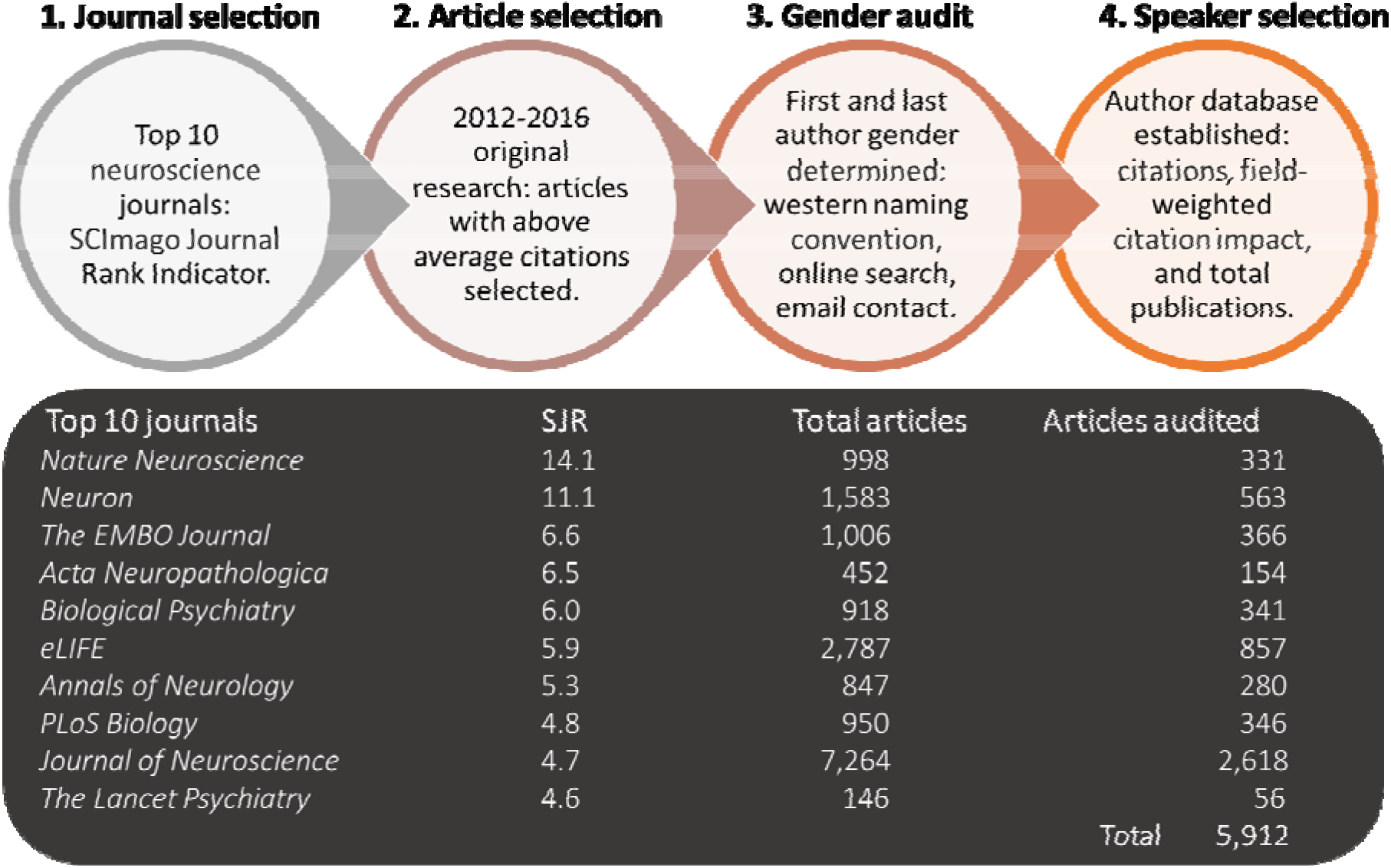
Selection procedure for creating a database for speaker selection.

### Journal Selection

Neuroscience journals were ranked using the SJR indicator system and Web of Science. The top ten journals comprising ≥50% original research articles were selected for auditing (see Figure 1). (Note: *Molecular Psychiatry* was excluded because more than 60% of publications reported authors’ initials only, making gender determination very difficult).

### Article Selection

Total citations and average citations per year were calculated for each original research article in the selected journals (citations from 2012-2016 for all journals except *Lancet Psychiatry*, for which citation data were only available from 2014-2016). Articles were selected for the author gender audit if their total citation count was greater than the average total citations for the journal in which the article was published.

### Gender identification

The gender of first and last authors of the selected articles was determined to be a man, woman, or unknown (last author was selected because it typically represents the senior author in neuroscience). Gender determination (using western naming convention) was completed independently by two investigators, and then cross-referenced. If gender could not be determined using this method, or the name was indeterminate or androgynous, an electronic search was conducted using institutional and academic networking websites: gender was determined if the online resources included the author’s name, photo (with clear gender identification) and either a reference to the article or the author’s affiliation (listed in the article). If gender of first or senior authors could not be determined using either of these methods (6.9%), the corresponding author was emailed to request gender identity information (email response rate: 20%). (In total, the gender of 163 authors could not be determined.)

### Quality metrics

The weighted total citations (2012-2016) were obtained by dividing the total citation counts for each paper by the number of years since its publication. The FWCI and their total number of career publications were obtained for the top 100 first and last authors (note: if an author did not have an identifiable FWCI using SciVal they were not included in the database.).

## Results

The 100 articles with the highest weighted total citations were used to create a database of potential speakers, including lists for first author and last author. The rank order within these lists was then adjusted based on FWCI. (All of the data for the top-100 first and senior authors are available supplementary material: ‘speaker database’.)

Critically, FWCI did not differ between men and women for either first or last authors (*p* > 0.49, Cohen’s *d* < 0.15, Bayes Factor, BF_10_ < 0.29), indicating no significant difference in the impact of research between men and women irrespective of career stage. Figure 2 shows the gender breakdown of authors in the top-100 database for FWCI and total publications. In the speaker database, 32% of first authors and 21% of last authors were women.

**Figure 2.**
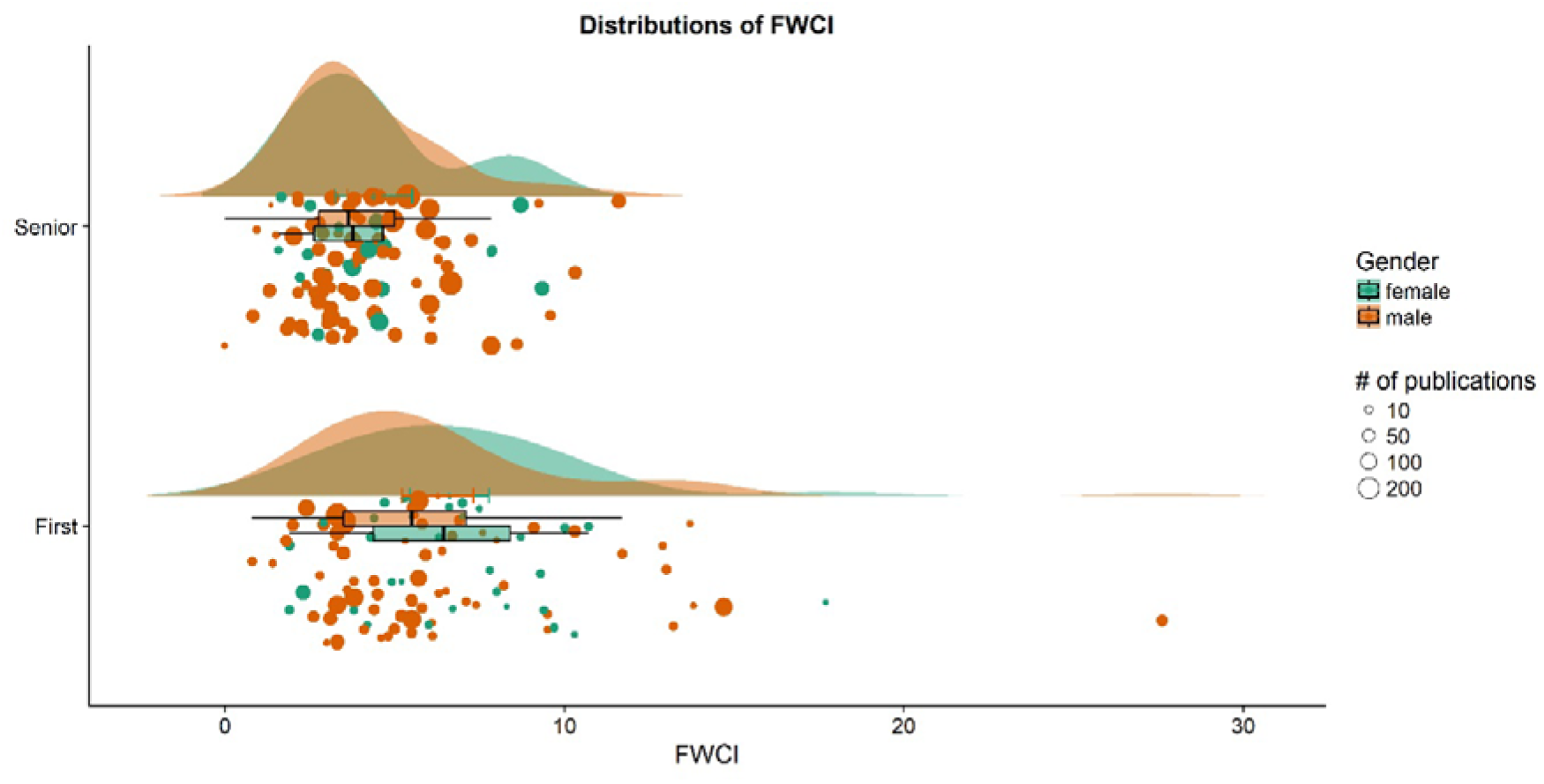
Raincloud plots of field weighted citation impact for women (green) and men (orange) first (lower plot) and senior (upper plot) authors in the top-100 database; each circle represents one author, with the size of the circle reflecting the total number of publications (see legend).

## Discussion

The data-driven approach presented here enables gender balanced speaker selection for conference programs based on scientific impact. Importantly, this approach, which assures the scientific quality of the speakers, can be used with gender policy to achieve gender parity in conference programs because the current results showed that scientific impact does not differ between men and women in the potential speaker database.

This data-driven approach to speaker selection takes an important step in addressing the complex issue of gender disparity in STEM, and extends beyond tools that are already available by identifying individuals as potential speakers and overcoming the criticism that selection based on policies is not quality-based. For example, online calculators can provide estimates of equitable gender representation, and equity and diversity policies can prescribe equitable gender representation, but neither provide any information regarding *who* to invite to deliver presentations. The current approach purposefully includes established metrics of scientific quality—that are frequently used by researchers, hiring committees, and funding bodies—to improve on existing approaches to ensure high quality speaker selection. Furthermore, the combination of metrics used in the approach presented here provides a database of potential speakers with a *recent* and *relevant* high-quality publication, whilst ensuring some stability in terms of career research performance.

Establishing the database of potential speakers using the current approach is largely automated (e.g., the exportation of publications, citations, FWCI), and the ranking of authors can be performed with simple code. For the current study, this process was neither arduous nor time-consuming. The identification of gender in the current study was somewhat time-consuming, however, this process could be automated if publishers required, and published, gender identity data: we urge journals to collect these data. In the current study, the broad discipline of neuroscience was the exemplar; using keywords and the selection of specialist journals, would make it possible to establish a database of potential speakers for a conference in a different discipline or for a focussed symposium within a conference. Indeed, we have previously shown that the proposed approach would be effective in the sub-discipline of brain stimulation (10). The data from the current study are available online, and we recommend that conference program committees use these data (together with gender policy), as well as continue to collect data, to reduce gender disparity in invited speaker programs.

It is important to acknowledge some limitations with the approach. First, achieving gender parity is not equal to achieving diversity and inclusion: the approach presented here should be extended to ensure representation of minority groups in conference programs. For example, expanding the approach to include geographical location, ethnicity, and career stage information would provide an opportunity to increase representation of minority groups in STEM. Second, the current approach relies on citations of publications in high impact journals. Evidence suggests that women submit fewer manuscripts than men to high quality journals, have fewer manuscripts accepted for publication in high quality journals, and that publications with a senior author who is a woman are cited less than publications with a senior author who is a man (e.g. 11). Therefore, although the approach presented here is data-driven, the data themselves are likely to be affected by biases that negatively affect women (1, 12). We strongly recommend that the approach presented here should be continually refined to include the most reliable and well-accepted quality metrics for STEM researchers.

It is worth noting that the underlying causes of gender disparity in STEM are not fully understood. Whilst beyond the scope of the current project, a growing literature strongly suggests that the persistence of gender disparity in STEM is due, at least in part, to implicit bias (13): the covert attitudes that influence our understanding, actions, and decisions in an unconscious manner. Evidence suggests that implicit gender bias in science negatively affects outcomes for women in terms of hiring, promotion, funding, and invitations for conference presentations, and editorial roles (e.g. 11, 14, 15); although note other studies suggest little bias against women (e.g. 16). The approach presented here minimizes implicit bias in speaker selection: given the strong evidence suggesting that implicit gender bias negatively affects women’s performance on the metrics used in this approach, we believe that the target for gender representation in conference program should be parity (not base rates, which are likely under-estimated (7)).

The benefits of the current approach are twofold. First, it provides a data-driven method for selecting invited speakers (senior and early career researchers), which can have an immediate effect on reducing gender disparity at scientific conferences. Second, establishing a database of high quality researchers based on metrics of scientific impact provides convincing evidence of parity in scientific quality between men and women at the highest level. These benefits should, in turn, lead to a positive spiral in which invited speaking opportunities for women facilitate career development through recognition of high-quality research, providing greater opportunity for collaborative outreach, which will increase likelihood of academic promotion and leadership for women within STEM. In light of the strengths and limitations of the current approach, we argue strongly that a combination of approaches will be most effective at reducing the persistent gender disparity, as well as increasing diversity in STEM more generally.

## Supporting information

speaker database

## Acknowledgments

We would like to thank Brigid Bolton, Ellika Carson, Courtney McAuliffe, Chelsea Moran, and Tayla Stucke for their assistance with this project. We would like to thank www.biaswatchneuro.com for providing data on invited speaker programs at neuroscience conferences.

## Author contributions

AMV, MRH, and HF conceived the idea, designed the study, interpreted results, and revised the manuscript; HF analysed data; AMV drafted the manuscript.

## Data and materials availability

all data is available in the manuscript or the supplementary materials.

Supplementary material: ‘speaker database’.

